# The Hippo pathway regulates density-dependent proliferation of iPSC-derived cardiac myocytes

**DOI:** 10.1101/2021.04.12.439529

**Authors:** Abigail C. Neininger, Xiaozhaun Dai, Qi Liu, Dylan T. Burnette

## Abstract

Inducing cardiac myocytes to proliferate is considered a potential therapy to target heart disease, however, modulating cardiac myocyte proliferation has proven to be a technical challenge. The Hippo pathway is a kinase signaling cascade that regulates cell proliferation during the growth of the heart. Inhibition of the Hippo pathway increases the activation of the transcription factors YAP/TAZ, which translocate to the nucleus and upregulate transcription of pro-proliferative genes. The Hippo pathway regulates the proliferation of cancer cells, pluripotent stem cells, and epithelial cells through a cell-cell contact-dependent manner, however it is unclear if cell density-dependent cell proliferation is a consistent feature in cardiac myocytes. Here, we used cultured <u>h</u>uman iPSC-derived <u>c</u>ardiac <u>m</u>yocytes (hiCMs) as a model system to investigate this concept. hiCMs have a comparable transcriptome to the immature cardiac myocytes that proliferate during heart development *in vivo*. Our data indicate that a dense syncytium of hiCMs can regain cell cycle activity and YAP expression and activity when plated sparsely or when density is reduced through wounding. We found that combining two small molecules, XMU-MP-1 and S1P, increased YAP activity and further enhanced proliferation of low-density hiCMs. Importantly, these compounds had no effect on hiCMs within a dense syncytium. These data add to a growing body of literature that link the Hippo pathway regulation with cardiac myocyte proliferation and demonstrate that regulation is restricted to cells with reduced contact inhibition.

## INTRODUCTION

Cardiac myocytes (CMs) are the cells of the heart that generate contractile force. The mechanisms controlling CM-proliferation are critical during both heart development and disease (Ponnusamy et al., 2017; Porrello et al., 2011). While adult CMs are non-mitotic, CMs in the developing heart proliferate to drive hyperplastic growth (Laflamme and Murry, 2011; Li et al., 1996; Walsh et al., 2010). The Hippo pathway has been shown to regulate proliferation during development (Heallen et al., 2011; Xin et al., 2013). The Hippo pathway is a kinase cascade that, when active, negatively regulates the transcriptional coactivators YAP and TAZ. In the absence of inhibition, YAP and TAZ translocate to the nucleus to activate the transcription of pro-proliferative genes (Dong et al., 2007; He et al., 2015; Zhao et al., 2010). In addition, the Hippo pathway regulates contact-dependent cell proliferation in epithelial cells, cancer cells, and pluripotent stem cells (Gumbiner and Kim, 2014; Kim et al., 2011; Totaro et al., 2017; Zhao et al., 2007). In these studies, low density cells with fewer cell-cell contacts have increased YAP nuclear localization, while high density cells have more inactive YAP in the cytosol (Schlegelmilch et al., 2011; Varelas, 2014). This evidence supports the notion that cell proliferation is not entirely regulated by paracrine mechanisms (e.g., by growth factors) and that proliferation is in part regulated spatially by density-dependent mechanisms (Kim et al., 2009). We asked if the Hippo pathway is involved in non-paracrine mechanisms such as contact inhibition in the heart.

The Hippo pathway is required for cardiac development in mice (von Gise et al., 2012; Wang et al., 2018). Cardiac-specific knockout of YAP is lethal in mice and leads to thin ventricular walls and several other structural abnormalities (Xin et al., 2013). A YAP mutant that blocks the interaction between YAP and its co-transcription factor TEAD also led to less proliferation in the developing mouse heart (von Gise et al., 2012). In addition, during the first post-natal week, the mouse heart retains the ability to fully regenerate after an infarction. YAP conditional knockout mice lose the ability to regenerate (Xin et al., 2013). Interestingly, constitutively active YAP can induce CM proliferation in the adult mouse heart, as can a mouse with a conditional knockout of Salv1, an upstream Hippo pathway component (Heallen et al., 2011). Taken together, these studies solidify the Hippo pathway as a key signaling cascade for CM proliferation.

We sought a model system to explore the role of the Hippo pathway in cell density-dependent regulation of CM proliferation. It is difficult to study contact inhibition *in vivo*. Controlling cell density is not possible. For example, acute mechanical perturbations followed by long-term live cell imaging are currently beyond available technologies. Therefore, we turned to *in vitro* models. It has previously been shown that plating low density human ESC-derived CMs induces them to enter the cell cycle (Park et al., 2018). This was shown using flow cytometry to measure the expression of three cell cycle markers: BrdU, YAP, and cyclin D1. Furthermore, our previous work showed that <u>h</u>uman iPSC-derived <u>CMs</u> (hiCMs) are more proliferative when plated sparsely than when in a monolayer based on cell counting using high-content microscopy (Neininger et al., 2019). Whether the Hippo pathway regulates the apparent difference in CM-proliferation capacity in dense and sparse populations remains to be elucidated. This led us to probe whether the Hippo pathway is involved in regulating the proliferative capacity of CMs in various densities or if YAP is activated in these sparse populations by another mechanism. We have chosen hiCMs as our model system to study this phenomenon, as they are transcriptionally immature—similar to fetal or neonatal CMs *in vivo*—and have a slight basal proliferative capacity (Uosaki et al., 2015; Sharma et al., 2018). Here, we show that reducing hiCM-density by either a scratch assay or by sparse plating increases nuclear YAP and CM-proliferation, and that this proliferative capacity can be further enhanced by combinatorial pharmacological perturbation of the Hippo pathway.

## RESULTS

### Reducing hiCM density in a scratch assay increases proliferative capacity

We first investigated whether a reduction in hiCM-density resulted in increased proliferation and/or any changes in Hippo pathway signaling. The hiCMs we used were purchased pre-differentiated and we confirmed the company’s claim of a lack of fibroblast contamination (See Materials and Methods) (Neininger et al., 2019). The lack of non-muscle cell types is key for interpretation of the data sets measuring proliferation. In addition, hiCMs can be cultured as a dense monolayer in which every cell has the potential to be experiencing contact inhibition.

One well-established method to relieve contact inhibition in culture is to scratch a confluent monolayer of cells. (Cory, 2011). Therefore, we used a micropipette tip to scratch a confluent monolayer of hiCMs (Figure 1A-B). We hypothesized that, as in epithelial cells, YAP would translocate to the nucleus of hiCMs near the scratch, as these hiCMs should have reduced contact inhibition compared to those far from the scratch. To test this, we localized YAP using immunofluorescence 48 hours post-scratch. We noticed that hiCMs near the edge of the scratch appeared to have primarily nuclear YAP (Figure 1C). A proxy for YAP activity is to measure the nuclear accumulation of YAP (Hansen et al., 2015). Therefore, we acquired higher magnification images to measure the ratio of nuclear YAP to total YAP. We used β-catenin to mark adherens junctions between myocytes in order to delineate separate cells, and measured nuclear YAP and total YAP using fluorescence localization. We found that hiCMs near the scratch had a higher nuclear YAP to total YAP ratio than either hiCMs farther from the scratch on the same plate or hiCMs in an unscratched monolayer (Figure 1C-D). Interestingly, this appeared to be an exponentially decaying relationship, where the nuclear YAP to total YAP ratio decreased rapidly as a function of distance from the scratch.

**Figure 1:**
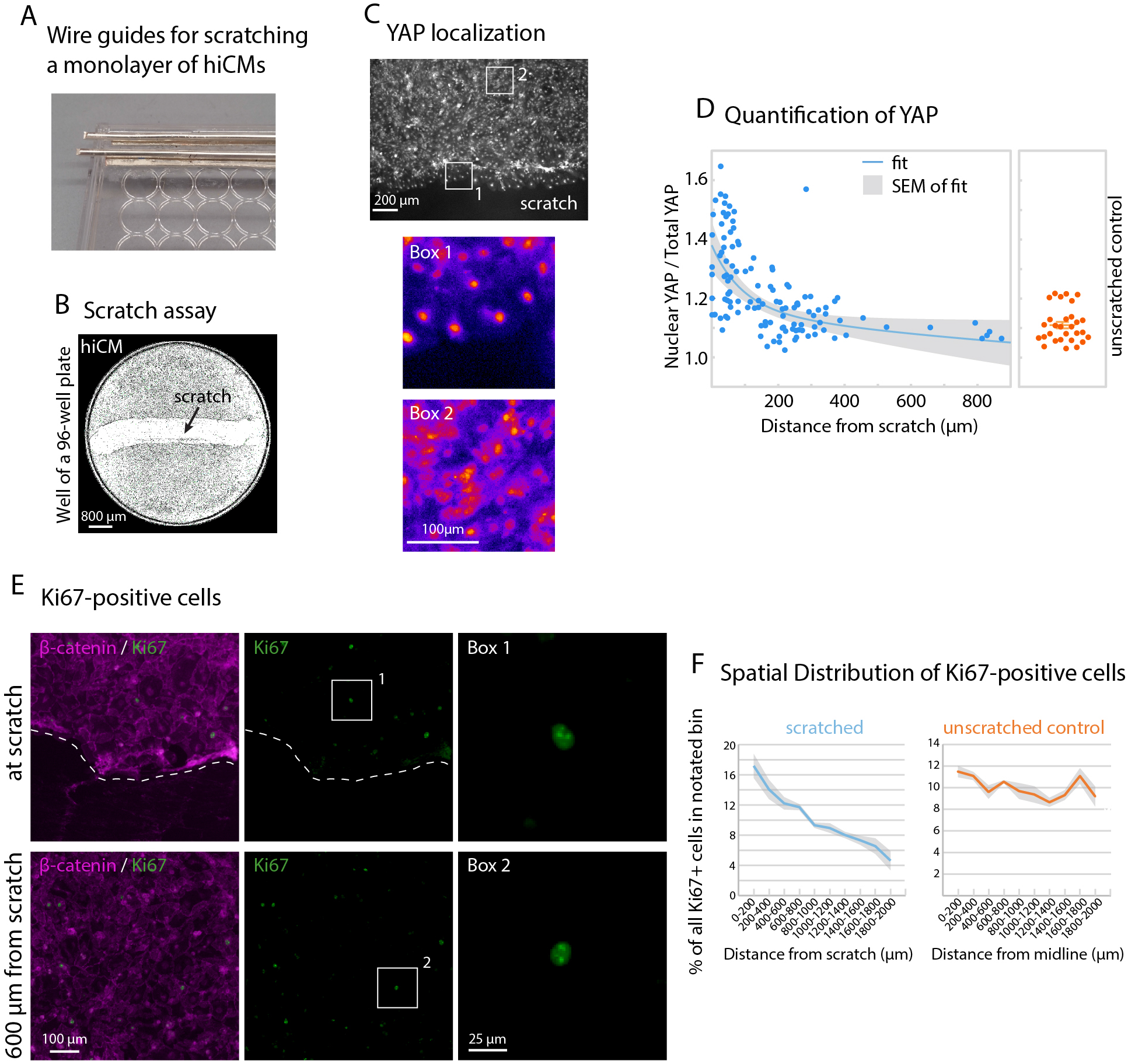
Reducing hiCM density by a scratch assay increases proliferative capacity. A) Wire guide for scratching a monolayer of hiCMs, using a lid of a 96-well plate and a Dremel tool to drill through the plastic lid. Wire guides are composed of a bottom layer of flattened 21-gauge wire glued to the lid and a top layer of cylindrical wire to guide the pipette tip. B) 4X whole-well phase-contrast image of hiCMs in a well of a 96-well plate, 48 hours post-scratch. C) YAP immunolocalization near the scratch (box 1) and far from the scratched (box 2) using 20X widefield microscopy. D) Quantification of Nuclear YAP divided by total YAP (n= 79 cells from 5 independent experiments). Unscratched control (n= 29 cells from 3 independent experiments). E) Ki67 and β-catenin in a scratched plate 48 hours post-scratch using 20X. F) Spatial distribution of Ki67-positive hiCMs in a scratched well or in an unscratched control well 48 hours post-scratch. ***p<0.0001, chi-square = 80.434 with a two by five contingency table comparing distributions of Ki67 in different regions within 2 mm of the scratch or center of the midline of the well.

We next wanted to know if the increase in nuclear YAP correlated with an increase in hiCMs entering the cell cycle. One standard way to identify cycling cells is to localize Ki67 using immunofluorescence, as Ki67 is localized in the nuclei of cycling cells (Gerdes et al., 1984; Miller et al., 2018). We found that in a monolayer of hiCMs, Ki67-positive cells were evenly distributed across the well (Figure 1E-F). In contrast, for the scratched cells, there was a skewed distribution, in which more Ki67-positive nuclei were near the scratch (Figure 1E-F). Taken together, these data indicate that the relief of contact inhibition by a scratch assay increases the amount of YAP in the nuclei of hiCMs and promotes their entry into the cell cycle. However, it is worth noting that a scratch assay is inherently asymmetric as only hiCMs near a scratch respond, limiting measurements of responses on a population level.

### Reducing hiCM density by sparse plating increases proliferative capacity

We next wanted to identify an assay that could model a uniform reduction in contact inhibition across an entire population of hiCMs. A previous study used flow cytometry and Western blotting to show that embryonic stem cell-derived cardiac myocytes begin cycling when they are sparsely plated (Park et al., 2018). Thus, we plated hiCMs as a monolayer or plated them sparsely. Not surprisingly, a monolayer of hiCMs (∼1088 hiCMs/mm^2^) have a low nuclear YAP/total YAP ratio (Figure 2A-B). On the other hand, when cells are plated at an 8-fold dilution (∼136 hiCMs/mm^2^), the nuclear YAP/total YAP ratio increases almost 2-fold (Figure 2A-B).

**Figure 2:**
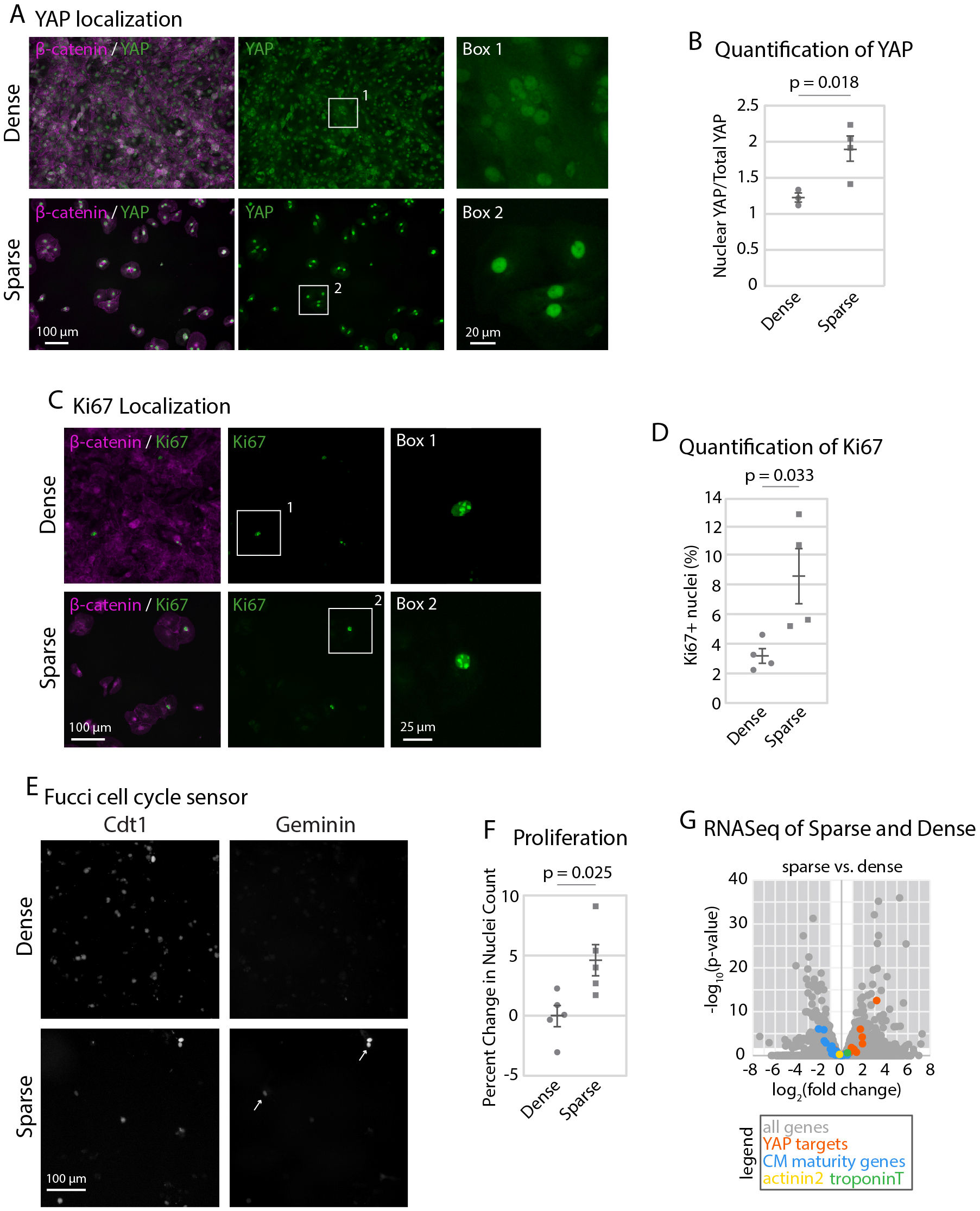
Reducing hiCM density by sparse plating increases proliferative capacity. A) YAP and β-catenin in dense and sparse hiCMs using 20X widefield fluorescence microscopy. B) Quantification of nuclear/total YAP ratio. Significance determined by paired two-tailed Student’s t test. (N = 23 dense cells and 27 sparse cells from 3 independent experiments) C) Ki67 and β-catenin immunolocalization in dense and sparse hiCMs using 20X widefield fluorescence microscopy. D) Percent of Ki67-positive nuclei in dense (N= 8046 hiCMs from 4 experiments) and sparse (N= 1716 cells from 3 experiments) hiCMs. Significance determined by an unpaired two-tailed Student’s t test E) Fucci cell cycle probe transduced into dense or sparse hiCMs. Arrows shows S-M phase. F) Proliferation of dense or sparse hiCMs quantified by counting nuclei fold change over 48 hours (N= 281250 cells from 5 independent experiments). Significance determined by unpaired two-tailed Student’s t test. G) RNASeq Volcano plot of genes significantly changed in sparse versus dense hiCMs. (N=300000 hiCMs per treatment from 3 independent experiments). A negative fold change represents a downregulation in sparse hiCMs. A positive fold change represents an upregulation in sparse hiCMs.

This increase in nuclear YAP in sparsely-plated hiCMs suggested an increase in proliferative capacity. To test this, we asked if sparse plating caused entry into the cell cycle using Ki67 localization. We localized Ki67 in dense or sparse hiCMs and found an increase in Ki67-positive nuclei, indicating a near doubling in cycling hiCMs (Figure 2C-D). We similarly used the Fucci probe system and found that the majority of dense hiCMs were not cycling, whereas sparse hiCMs had increased cycling cells (Figure 2E). To directly test if sparse-plating induced proliferation in hiCMs, we counted the number of hiCMs over time. We found that while dense control hiCMs essentially remained the same over 48 hours, there was a ∼5% increase in the number of sparsely-plated hiCMs (Figure 2F). Taken together, our data indicates that sparse-plating induces hiCMs to enter the cell cycle and also divide. This led us to explore the underlying cause of sparsely-plated hiCMs to enter the cell cycle.

It has been postulated that CM division is preceded by dedifferentiation into a more immature phenotype (D’Uva et al., 2015; Wang et al., 2017). Thus, we employed RNASeq analysis to examine whether the proliferating sparsely-plated hiCMs were as mature as hiCMs in a monolayer. We found CM-maturity genes (Uosaki et al., 2015, Table S1) were modestly downregulated in sparse hiCMs (Figure 2G). However, expression of CM-identity genes α-actinin 2 and troponinT did not change (Figure 2G, Table S3). Thus, sparsely-plated hiCMs are less mature than densely-plated hiCMs, but they remain CMs.

We next wanted to test if the increase in nuclear YAP ratio in sparsely-plated hiCMs was correlated with higher YAP target gene expression. Thus, we curated a list of YAP target genes from the published literature (Kim et al., 2018; Lee et al., 2016; Venkataramani et al., 2018, Table S2). We then compared the expression of these genes between densely-plated and sparsely-plated hiCMs. 32% of YAP target genes had upregulated expression in sparsely-plated hiCMs compared to dense, while 68% did not change, and 0% had decreased expression. In conclusion, plating hiCMs sparsely increases YAP nuclear localization, YAP target gene expression, and ultimately, hiCM proliferation compared to densely-plated hiCMs.

### Pharmacological perturbation of the Hippo pathway affects cardiac myocyte proliferation

A loss of contact inhibition not only increased hiCM proliferation but also the nuclear YAP/total YAP ratio, consistent with the hypothesis that the Hippo pathway is involved in density-dependent hiCM proliferation. We next asked if small molecule inhibition of the Hippo pathway would further increase the proliferative potential of hiCMs plated at a low density. We turned to an MST1/2 inhibitor, XMU-MP-1, and a bioactive lipid that activates YAP, sphingosine-1-phosphate (S1P) (Fan et al., 2016; Miller et al., 2012; Sharma et al., 2018) (Figure 3A). Both of these compounds have been shown to upregulate YAP target genes in human embryonic stem cells, mouse liver cells and human hepatoma cells (Fan et al., 2016; Venkataramani et al., 2018). Furthermore, S1P increases Ki67 in densely plated hiCMs but not cell count when combined with lysophosphatidic acid (Sharma et al., 2018).

**Figure 3:**
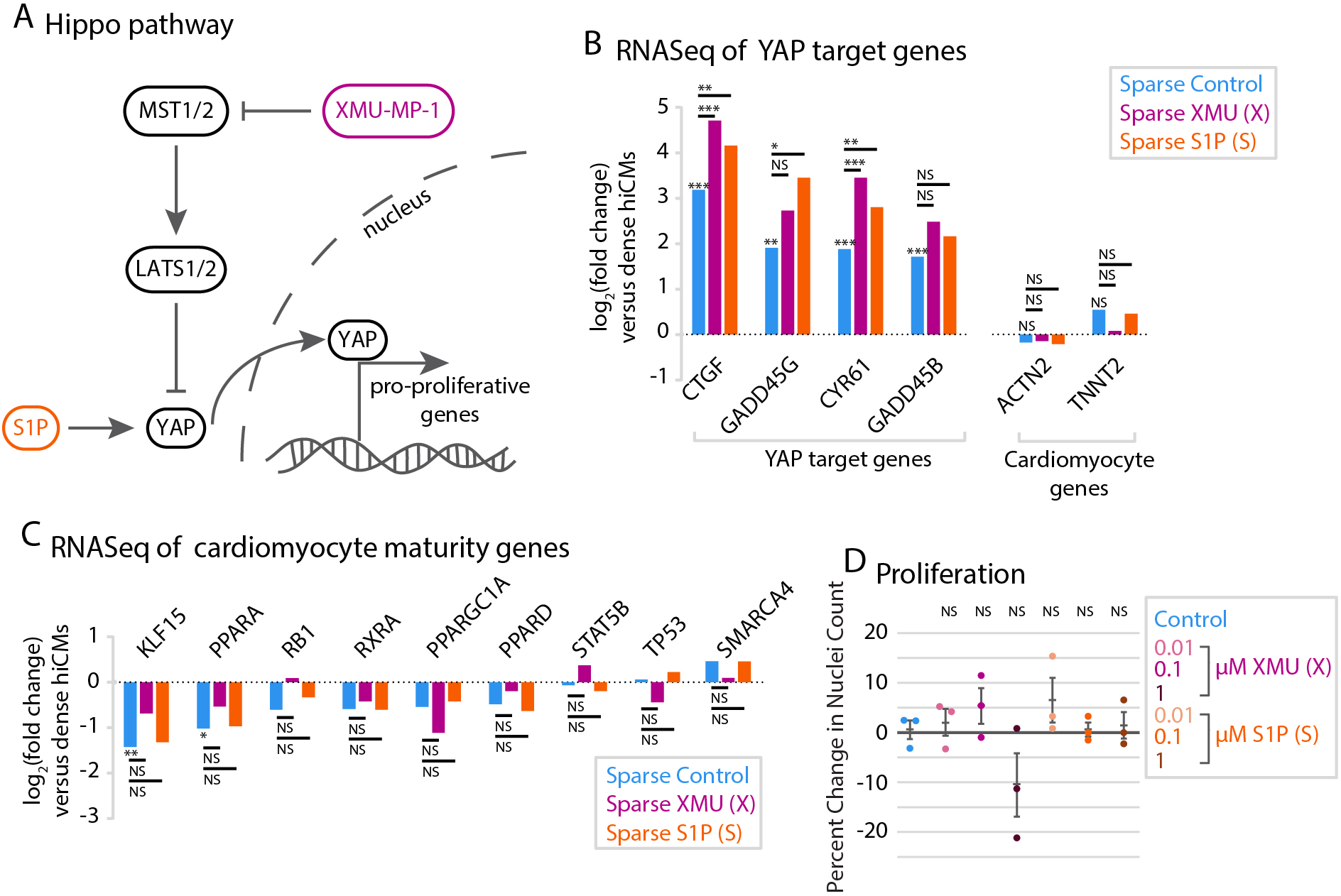
Individual pharmacological perturbation of the Hippo pathway increases YAP target genes but does not affect proliferation. A) Schematic of the Hippo pathway. B) RNASeq data of YAP target genes in sparse cells with treatment of 1 µM XMU-MP-1 or 1 µM S1P. A negative fold change represents a downregulation in treated cells, whereas a positive fold change represents an upregulation in treated cells. N=300000 cells per treatment, over 3 independent experiments and RNA preparations. Right: cardiac myocyte-specific genes do not change upon small molecule treatment. *: p < 0.05, **: p < 0.01, ***, p < 0.001. All p-values and gene names are available in Tables S1-S3. Differentially expressed genes were determined using the criteria of fold change >= 2 and FDR <= 0.05. C) RNASeq data of CM-maturity genes as in B. D) Quantification of proliferation of control, XMU-MP-1-, and S1P-treated cells by measuring the percent change in nuclei over 48 hours post-treatment. Significance determined by one-way ANOVA with a Dunnett’s post-hoc test corrected for multiple testing.

We used RNASeq analysis to test if XMU-MP-1 or S1P upregulate YAP/TAZ target genes in hiCMs, and found that YAP/TAZ target genes were indeed significantly upregulated beyond that of sparse-plating alone (Figure 3B, Table 1). We then confirmed that the hiCMs were still expressing CM specific markers (Figure 3B, indicating that XMU-MP-1 and S1P further increase YAP target gene expression in hiCMs. This increase in YAP target gene expression suggests that the compound-treated hiCMs are less mature than untreated sparsely-plated hiCMs. We found that S1P treatment did not lower the expression of maturity genes compared to untreated hiCMs, whereas 4.2% of maturity genes were expressed lower after XMU-MP-1 treatment (Figure 3C, Table 2). Taken together, our data indicates that XMU-MP-1 or S1P treatment causes an upregulation of YAP/TAZ target genes with minor effects on maturity of hiCMs.

Despite an upregulation in YAP/TAZ target genes, we found that neither XMU-MP-1 nor S1P alone increased proliferation by hiCM count (Figure 3D). The result that XMU-MP-1 or S1P failed to induce proliferation was not surprising, as previous studies have postulated that combinations of multiple genetic or pharmacological factors are required to induce cardiac myocyte proliferation both *in vitro* and *in vivo* (Mills et al., 2019; Mohamed et al., 2018). Therefore, we performed a combinatorial screen to determine if combining XMU-MP-1 and S1P induced proliferation of hiCMs. We started by testing which combination of concentrations was most effective to induce hiCMs to reenter the cell cycle using BrdU incorporation (Figure 4A). We found that adding 0.1 µM of each compound to hiCMs significantly increased the percent of BrdU-positive hiCMs when sparsely-plated (Figure 4B), but not in a dense syncytium (Figure 4D). We then found that this combination significantly increased hiCM-proliferation by counting nuclei over time (Figure 4C). We found that using 0.1 µM of both XMU-MP-1 and S1P also decreased phospho-YAP, which indicated a reduction in the inactivation of YAP (Figure 4E-F).

**Figure 4:**
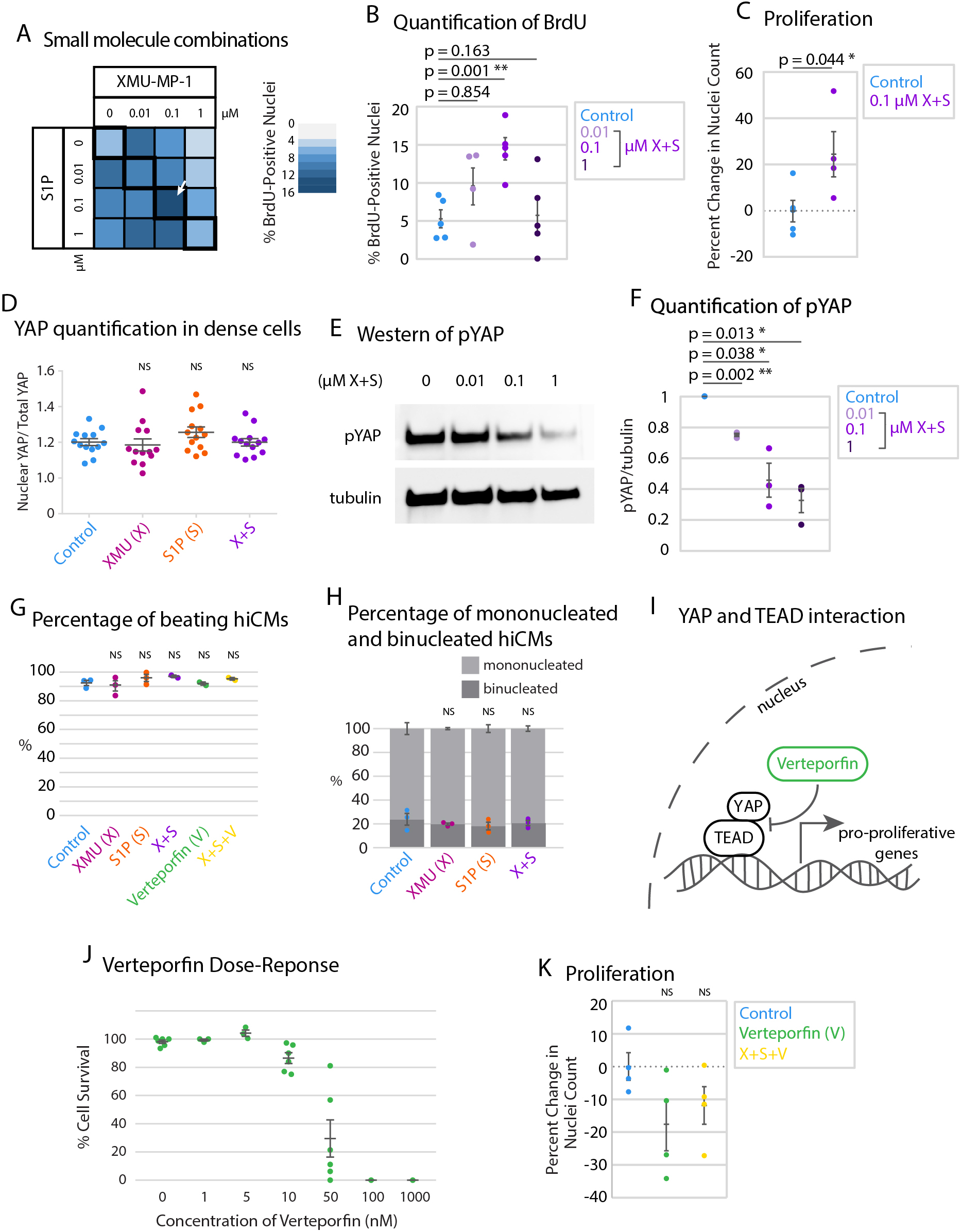
Combination of XMU-MP-1 and S1P increases cardiac myocyte proliferation. A) Heat map showing percentage of BrdU-positive nuclei in a preliminary screen of combinatorial treatment of hiCMs with XMU-MP-1 and S1P. N = 8393 cells from 5 independent experiments. B) BrdU+ datapoints from black outlined diagonal center boxes from heat map. N = 2229 cells from 5 independent experiments. C) Proliferation of control hiCMs or 0.1 µM XMU-MP-1 and 0.1 µM S1P-treated cells measured by counting nuclei over 48 hours. N = 17318 DMSO-treated cells and 16234 XS-treated cells from 4 independent experiments. Significance determined by two-tailed paired Student’s t-test. D) Western blot of hiCMs treated with various concentrations of both XMU-MP-1 and S1P. pYAP: phosphorylated YAP. E) XMU-MP-1 and/or S1P does not increase nuclear YAP/total YAP ratio in dense hiCMs. F) pYAP amount in hiCMs measured by western blot normalized to tubulin (N= 3 independent experiments). G) The compounds XMU-MP-1, S1P, and Verteporfin do not affect the percentage of beating hiCMs at the doses given (1 µM for XMU-MP-1 and S1P, 0.1 µM for XMU-MP-1 and S1P combined, and 10 nM Verteporfin. In last row, 0.1 µM of XMU-MP-1 and S1P were combined with 10 nM Verteporfin). H) The compounds XMU-MP-1 and S1P do not affect the percentage of binucleated and mononucleated hiCMs at the same doses as Figure 4G. I) Schematic of YAP’s downstream transcriptional regulation and interaction with TEAD. J) Verteporfin dose response to detect which concentration of Verteporfin inhibited hiCM proliferation, but did not lead to massive cell death. K) Proliferation of control hiCMs compared to 0.01 µM Verteporfin, or 0.01 µM Verteporfin, 0.1 µM S1P, and 0.1 µM XMU-MP-1-treated hiCMs. N = 1257 cells from 4 independent experiments. Significance determined by one-way ANOVA with a Dunnett’s post-hoc test correcting for multiple testing for panels B, D, F, G, H, and K.

It is important to confirm that these proliferative hiCMs are functionally beating upon treatment of XMU-MP-1 and S1P. Therefore, we measured the percentage of beating hiCMs upon sparse plating and treatment of XMU-MP-1, S1P, or both, and found no significant difference (Figure 4G). Further, it is important to consider that both cytokinetic cell division and binucleation are potential outcomes of increasing overall proliferation of cells. We measured the percentage of binucleated hiCMs upon sparse plating and treatment of XMU-MP-1, S1P, or both, and found no significant difference (Figure 4H). In this case, it is presumed that proliferation appears to be induced overall, but the balance between the potential outcomes of cell division and binucleation are not changed, and both outcomes are occurring. These results agree with past research inducing hiCMs to divide, showing both binucleation and cell division as outcomes (Neininger et al., 2019).

Finally, to further confirm that XMU-MP-1 and S1P were acting through the Hippo pathway, we turned to Verteporfin, which inhibits the interaction of YAP with transcription factor TEAD (Liu-Chittenden et al., 2012) (Figure 4I). For this experiment we chose 0.01 µM of Verteporfin, which is the highest concentration that did not result in the death of hiCMs (Figure 4J). We found that Verteporfin masked the effect of the combination of XMU-MP-1 and S1P combination on proliferation (Figure 4K) without affecting the percentage of beating hiCMs (Figure 4G). This result further suggests that the increase in proliferation of hiCMs with XMU-MP-1 combined with S1P is through the Hippo pathway.

## DISCUSSION

Taken together, we show that the density at which hiCMs are plated affects the cells proliferative capacity, and that this proliferative capacity is regulated at least in part by the Hippo pathway. Further, we modulate the proliferative capacity by scratching a monolayer of cells to induce cells at the scratch periphery to divide, and induce division of sparsely-plated hiCMs by dual inhibition of the Hippo pathway. These experiments were completed using hiCMs, a model system of developmentally immature cells that both divide and binucleate, providing an ideal model for studying cardiomyocyte division. This is an alternative approach to using isolated primary cardiomyocytes from adult rats or mice, which are unsuitable to the experiments we use here. While primary cardiomyocytes are powerful tools, creating dense monolayer is technically challenging. In addition, primary cardiomyocytes only last a few days in culture and are more representative of modeling cell death (Peter et al., 2016). Therefore, it is difficult to do contact inhibition studies and proliferative studies in these cell types. As we show here, hiCMs are well suited for these types of studies (Neininger et al., 2019).

It is not surprising that XMU-MP-1 itself did not increase CM proliferation, as a recent study found that XMU-MP-1 in an intact heart did not increase Ki67 staining (Triastuti et al., 2019). Based on Ki67 staining, recent data has also suggested that inhibiting the Hippo pathway with a single compound in combination with perturbation of another cell cycle pathway (i.e., the Wnt pathway) could be an effective approach to increasing CM proliferation (Mills et al., 2019). Here, we show that it is possible to have a combinatorial effect on CM proliferation by only targeting the Hippo pathway. Modulation of the Hippo pathway at two different levels was required to get a proliferative response. This could be part of a new direction for combinatorial therapy, where multiple pathways are each modulated at multiple levels.

There are several other pathways that drive CM proliferation. A recent study suggests the neuregulin receptor, ERBB2, activates YAP in an ERK-dependent manner (Aharonov et al., 2020). The role of ERK signaling in cardiomyocyte proliferation and its crosstalk with the Hippo pathway would constitute an interesting future study. Furthermore, the signaling role that cell-cell adhesions and junctions themselves play in the cardiac regenerative response seems to merge on these pathways. Regenerating cardiac myocytes undergo junction dissolution, and it has been shown that displaced alpha-catenin activates YAP in epidermal cells (Schlegelmilch et al., 2011). Finally, increasing YAP and TAZ activity is likely to have benefits in the heart other than increasing cardiac myocyte proliferation. For example, YAP and TAZ in the epicardium was shown to induce recruitment of T-regulatory cells to the infarcted myocardium. Thus, YAP and TAZ may have the ability to regulate the adaptive immune response and decreasing post-infarct inflammation and myocardial fibrosis (Ramjee et al., 2017).

Finally, our work also has implications in the context of a myocardial infarction. In the developed heart, a myocardial infarction results in massive CM death and the remaining CMs do not proliferate (Pfeffer & Brunwald, 1990). After a myocardial infarct, there is a reduction in cell density at the infarct zone with an increase in cell cycling shown by Ki67 fluorescence (Meckert et al., 2005). However, it has been shown that the proliferative cells in the infarct region appear to mostly undergo polyploidization and endomitosis, indicating that there is a reduction in contact inhibition but there is also still a barrier to true cytokinetic proliferation in this region. If an ideal treatment for a cardiac infarction existed, it would only induce proliferation at the infarct site where cell-cell contact is reduced. Global induction of proliferation could induce undesired side effects (e.g. tumor formation). As such, targeting pathways which respond preferentially when contact inhibition is lost could be a useful therapeutic direction. Here, we show that targeting the Hippo pathway can increase proliferation in the context of a loss of contact inhibition.

## MATERIALS AND METHODS

### KEY RESOURCES

| REAGENT or RESOURCE | SOURCE | IDENTIFIER |
| --- | --- | --- |
| **Antibodies** |  |  |
| Mouse-anti-α-actinin-2 | Sigma | A7811 Clone: EA53 |
| Mouse anti-β-catenin | BD Biosciences | 610153 |
| Rabbit-anti-YAP | Cell Signaling Technologies | 14074T |
| Rabbit-anti-phosphoYAP | Cell Signaling Technologies | 13008S |
| Mouse-anti-α-tubulin | Sigma | B512 Clone: DM1α |
| Rabbit-anti-Ki67 | Cell Signaling Technologies | 9129S |
| Mouse-anti-BrdU | Cell Signaling Technologies | 5292 |
| Goat anti-mouse 488 | LifeTechnologies | A11001 |
| Goat anti-rabbit 488 | LifeTechnologies | A11034 |
| Goat anti-mouse 568 | LifeTechnologies | A11004 |
| Goat anti-rabbit 568 | LifeTechnologies | A11036 |
| Goat anti-mouse HRP | LifeTechnologies | 62-6520 |
| Goat anti-rabbit HRP | LifeTechnologies | 65-6120 |
| **Biological Samples** |  |  |
| Cardiac myocytes<sup>2</sup> | Cellular Dynamics | CMC-100-012-000.5 |
| Bovine Serum Albumin | RPI | A30075-100.0 |
| **Chemicals, Peptides, and Recombinant Proteins** |  |  |
| NucLight Rapid Red | Essen Biosciences | 4717 |
| Vectashield with DAPI | Vector | H-1200 |
| Cardiac myocyte Plating Medium | Cellular Dynamics | M1001 |
| Cardiac myocyte Maintenance Medium | Cellular Dynamics | M1003 |
| PBS, 10X, Ca<sup>2+</sup>/Mg<sup>2+</sup> free | Gibco | 70011-044 |
| Paraformaldehyde, 16% | Electron Microscopy Sciences | 15710 |
| PBS, 10X, with Ca<sup>2+</sup>/Mg<sup>2+</sup> | Corning | 46-013-CM |
| 0.5% Trypsin | Gibco | 15400-054 |
| 0.1% Gelatin | Sigma | ES-006-B |
| Dimethyl Sulfoxide | Sigma | 276855 |
| Triton X-100 | Fisher Scientific | BP151100 |
| Fibronectin | Corning | 354008 |
| BrdU | Abcam | Ab142567 |
| XMU-MP-1 | SelleckChem | S8834 |
| S1P | Apex Bio | B6707 |
| Methanol | Fisher Scientific | A452SK-1 |
| Verteporfin | Sigma | SML0534 |
| Premo FUCCI Cell Cycle Sensor | Invitrogen | P36238 |
| **Software and Algorithms** |  |  |
| FIJI/ImageJ | NIH, Open Source |  |
| Incucyte analysis software | Essen Biosciences |  |
| MATLAB | MathWorks |  |

#### Cell culture and chemicals

iPSC-derived human cardiac myocytes (hiCMs, Cellular Dynamics, Madison, WI) were seeded as per manufacturer’s instructions in cardiac myocyte plating medium (M1001, Cellular Dynamics, Madison, WI) in polystyrene 96-well cell culture plates coated in 0.1% sterile gelatin (ES-006-B, Sigma Aldrich, St. Louis, MO). Approximately 50,000 cells were seeded for dense plating and 6,250 cells for sparse plating (1:8), and were subsequently cultured in cardiac myocyte maintenance medium (M1003, Cellular Dynamics, Madison, WI). These cells are verified by Cellular Dynamics (Madison, WI), and each vial of cells is tested at the company for purity by cTNT+ cells using flow cytometry (over 99% pure), identity using SNP genotyping, mycoplasma testing by PCR, sterility testing by 21 CFR 610.12, MEA functionality testing by field potential duration and beating rate, and viability by trypan blue exclusion. We further characterized these cells in Neininger et al., 2019, in which we determined that fewer than 1 in 50,000 cells were actinin2-negative.

Cells were kept at 37°C and 5% CO_2_. For re-plating experiments, hiCMs were washed 2x with 100uL 1x PBS with no Ca2+/Mg2+ (PBS*, 70011-044, Gibco, Grand Island, NY). PBS* was completely removed from hiCMs and 40uL 0.1% Trypsin-EDTA with no phenol red (15400-054, Gibco, Grand Island, NY) was added to hiCMs and placed at 37°C for 2 minutes. Following incubation, culture dish was washed 3x with trypsin inside well, rotated 180 degrees, and washed another 3x. Trypsinization was then quenched by adding 120 µL of culture media and total cell mixture was placed into a 1.5mL Eppendorf tube. Cells were spun at 1000gs for 3 minutes, and supernatant was aspirated. Cells were then re-suspended in 200uL of culture media and plated into 2 wells, 100 uL each, on a standard polystyrene 96-well cell culture plate (3599, Corning, Corning, NY) previously coated with 10 ug/mL Fibronectin (#354008, Corning, Mannassas, VA) for 1 hour at 37°C.

XMU-MP-1 (S8334, Selleck Chemicals, Houston, TX) and Verteporfin (SML0534, Sigma Aldrich, St. Louis, MO) were reconstituted to 10 mM in DMSO. S1P (B6707, Apex Bio, Houston, TX) was reconstituted to 10 mM in methanol. When adding drugs to the cells, they were first diluted 1:10 in DMSO then to the appropriate concentration in cardiac myocyte maintenance medium. NucLight Rapid Red Reagent (4717, Essen BioScience, Ann Arbor, MI) was used at a 1:4000 dilution in cell culture media.

#### Fixation and immunostaining

Cells were fixed with 4% paraformaldehyde (PFA, 15710, Electron Microscopy Sciences, Hatfield, PA) diluted form 16% in PBS at room temperature for 20 min, then extracted for 5 min with 1% Triton X-100 (BP151100, Fischer Scientific, Suwanee, GA) and 4% PFA in PBS. Cells were washed three times in 1× PBS. After fixation, the following labeling procedures were used: for immunofluorescence experiments, cells were blocked in 10% bovine serum albumin (BSA) in PBS for 20 minutes. Primary antibodies were diluted in 10% BSA. All primary antibodies were used at 1:200 for 1 hour and 45 minutes at room temperature. Secondary antibodies were diluted in 10% BSA at 1:100 and centrifuged at 13,000 rpm for 2 min before use at room temperature for 1 hour.

Mouse anti-β-catenin was purchased from BD Biosciences (610153). Rabbit anti-Ki67 (9129S) and Rabbit anti-YAP (14074T) were purchased from Cellular Signaling Technologies (Danvers, MA). Mouse anti-actinin-2 (a7811) was purchased from Sigma Aldrich (St. Louis, MO). Rabbit anti-phosphoYAP (13008S) was purchased from Cellular Signaling Technologies (Danvers, MA). Mouse anti-BrdU (5292S) was purchased from Cellular Signaling Technologies (Danvers, MA). Alexa Fluor 488-goat anti-mouse (A11029), Alexa Fluor 488-goat anti-rabbit (A11034), Alexa Fluor 568-goat anti-rabbit (A11011), and Alexa Fluor 568-goat anti-mouse (A11004) antibodies were purchased from Life Technologies (Grand Island, NY).

BrdU (ab142567, abcam, Cambridge, UK) was reconstituted to 10 mM in water, then to 10 µM in cardiomyocyte maintenance medium and filtered through a 0.22 µm syringe filter. Cells were incubated with each compound and BrdU for 24 hours, then fixed and permeabilized as above. Then, to hydrolyze DNA, cells were incubated with 1 M HCl for an hour at room temperature, then neutralized with 0.1 M sodium borate (pH 8.5) for 20 minutes at room temperature. Next, cells were washed with PBS and immunostaining continued as usual.

#### Scratch assay

Scratches in 10 mm dishes were done by hand with a 10 µL pipette tip. Scratches in 96-well cell culture polystyrene plates were done using a 200 µL pipette tip and a custom plate lid produced by A.C.N. with a slot designed to guide the pipette tip straight across the well. Media was replaced with media containing NucLight Rapid Red Reagent immediately after scratching and every two days afterward until fixation at specified time points.

#### Western Blotting

Cell lysates were prepared by lysing 50,000 cardiac myocytes with CellLytic (Sigma, St. Louis, MO, #C2978) with 1% protease inhibitor cocktail (P8340, Sigma Aldrich, St. Louis, MO). Gel samples were prepared by mixing cell lysates with LDS sample buffer (Life Technologies, #NP0007) and Sample Reducing Buffer (Life Technologies, #NP00009) and boiled at 95°C for 5 minutes. Samples were resolved on Bolt 4-12% gradient Bis-Tris gels (Life Technologies, #NW04120BOX). Protein bands were blotted onto a nylon membrane (Perkin Elmer, Boston MA, NBA085C001EA) with western blotting filter paper (ThermoFisher Scientific, Rockford, IL, #84783). Blots were blocked using 5% nonfat dairy milk (Research Products International Corp, Mt. Prospect, IL, #M17200) in TBST (TBS: Corning, Mannassas, VA, #46-012-CM. Tween20: Sigma, St. Louis, MO, #P9416). Antibody incubations were also performed in 5% NFDM in TBST. Blots were developed using the Immobilon Chemiluminescence Kit (Millipore, #WBKLS0500).

#### Statistics and quantification

Total YAP was measured in FIJI (ImageJ, NIH, Bethesda, MD) by drawing a polygonal ROI around a cell using a 20x IncuCyte image stained for B-catenin to mark cell boundaries, and measuring average fluorescence intensity. Nuclear YAP was measured in FIJI by drawing a Bezier ROI around a nucleus using a 20x IncuCyte image stained for YAP and measuring average fluorescence intensity. Fit of nuclear YAP / total YAP graph in Figure 1D was determined using the curve fitting toolbox in MatLab (MathWorks, Natick, MA). Nuclear fold change was determined by thresholding live-cell whole-well 4x stitches with NucLight Rapid Red Reagent, a live-cell nuclear marker. Thresholding was done using the IncuCyte software (Essen Biosciences, Ann Arbor, MI) and a TopHat Background subtraction using a 20 µM rolling ball. Images were acquired and nuclei counted every hour for specified time points. Nuclei count at the final time point was divided by nuclei count at the first time point and normalized to the average nuclear fold change of cells in control conditions on the same plate.

Statistical significance of Figures 2D and 2F were determined by unpaired two-tailed Student’s t-tests performed in Excel. Statistical significance of Figures 2B and 4C were determined by paired two-tailed Student’s t-tests performed in Excel. Statistical significance of Figures 3D, 4B, D, F, G, H, and K were determined by one-way ANOVA with a Dunnett’s post-hoc test correcting for multiple testing when applicable performed in GraphPad Prism. Each experiment was performed a minimum of 3 times and the mean of the independent experiments and standard error of the mean (SEM) are displayed.

#### RNA Sequencing

Cell pellets of 100,000 cells were prepared and RNA was extracted using the RNeasy Mini Kit (Qiagen, #74104). Stranded mRNA (polyA-selected) library preparation was completed by the VANTAGE core at Vanderbilt University. Sequencing was done with an Illumina NovaSeq6000 (S4) PE150. Reads were mapped to reference genome using STAR and differential analyses were performed using DESeq2. Differentially expressed genes were determined using the criteria of fold change >=2 and FDR <= 0.05. YAP target genes and cardiac myocyte maturity genes were determined from literature and are listed below. Datasets will be uploaded to a public database. N=3 for each condition, in that each cell treatment and RNA preparation was done three separate times using three separate purchased vials of cells.

## ACKNOWLEDGEMENTS

We would like to thank the VANTAGE core at Vanderbilt University for RNA Sequencing, and James B. Hayes and Manuel Castro (Vanderbilt University) for careful reading and comments on the manuscript.

## AUTHOR CONTRIBUTIONS

A.C.N and D.T.B. conceived the project and wrote the manuscript. A.C.N designed, performed, and quantified experiments, and prepared figures. X.D. and Q.L. analyzed RNASeq data.

## FIGURES

**Table S1:** YAP Target Genes

| Gene | Sparse<br>(Dense)<br>FC | p-value | XMU<br>(Dense)<br>FC | p-value | S1P<br>(Dense)<br>FC | p-value | XMU<br>(Sparse)<br>FC | p-value | S1P<br>(Sparse)<br>FC | p-value |
| --- | --- | --- | --- | --- | --- | --- | --- | --- | --- | --- |
| CTGF | 9.01 | 2.89E-13 | 25.81 | 2.79E-22 | 17.62 | 8.25E-24 | 2.93 | 0.000134 | 1.97 | 0.007319 |
| GADD45G | 3.73 | 0.002072 | 6.56 | 0.000193 | 10.85 | 0.000115 | 1.79 | 0.605755 | 3.00 | 0.038005 |
| CYR61 | 3.66 | 5.95E-05 | 10.82 | 1.04E-13 | 6.92 | 6.18E-14 | 3.01 | 0.000925 | 1.91 | 0.020171 |
| GADD45B | 3.26 | 9.59E-07 | 5.54 | 1.41E-08 | 4.45 | 0.001261 | 1.74 | 0.126936 | 1.39 | 0.36164 |
| GLI2 | 1.53 | 0.393604 | 3.82 | 6.2E-22 | 2.24 | 0.185947 | 2.54 | 0.004467 | 1.47 | 0.316052 |
| CDKN2B | 2.54 | 0.175283 | 2.99 | 0.000286 | 2.57 | 0.384279 | 1.22 | 0.792722 | 1.01 | 0.983649 |
| MYC | 2.11 | 0.065524 | 3.30 | 9.61E-08 | 3.98 | 7.42E-08 | 1.60 | 0.275537 | 1.90 | 0.012205 |
| VIM | 1.85 | 0.026872 | 1.39 | 0.000517 | 1.68 | 0.038557 | 0.77 | 0.478305 | 0.92 | 0.7466 |
| TEAD4 | 1.87 | 0.015487 | 1.63 | 9.48E-06 | 1.68 | 0.023656 | 0.89 | 0.778883 | 0.91 | 0.679315 |
| GPATCH4 | 2.18 | 0.038921 | 0.78 | 0.115272 | 2.01 | 0.357642 | 0.37 | 0.002401 | 0.93 | 0.87144 |
| TXN | 1.69 | 0.183088 | 1.36 | 0.039324 | 1.96 | 0.299241 | 0.82 | 0.697661 | 1.18 | 0.672179 |
| CAV1 | 0.63 | 0.143092 | 0.33 | 4.8E-18 | 0.62 | 0.135195 | 0.53 | 0.004174 | 1.00 | 0.99217 |
| CAT | 0.81 | 0.520764 | 0.84 | 0.245518 | 0.73 | 0.551213 | 1.06 | 0.871611 | 0.90 | 0.656182 |
| TEAD1 | 1.08 | 0.904655 | 1.70 | 3.93E-06 | 1.31 | 0.694522 | 1.61 | 0.246832 | 1.21 | 0.571363 |
| AXL | 0.71 | 0.199238 | 0.62 | 0.000225 | 0.70 | 0.260978 | 0.88 | 0.69564 | 0.99 | 0.962445 |
| CCND1 | 1.21 | 0.480377 | 0.99 | 0.95182 | 1.15 | 0.630965 | 0.83 | 0.467474 | 0.96 | 0.791495 |
| ERBB4 | 0.82 | 0.630638 | 0.95 | 0.814267 | 1.01 | 0.995008 | 1.18 | 0.703516 | 1.22 | 0.533846 |
| WSB2 | 0.91 | 0.780978 | 0.81 | 0.033037 | 0.82 | 0.757739 | 0.90 | 0.726643 | 0.90 | 0.696159 |
| LMNB2 | 1.12 | 0.807049 | 0.97 | 0.900838 | 1.08 | 0.886014 | 0.89 | 0.761298 | 0.97 | 0.88176 |
| TNFAIP3 | 0.74 | 0.686538 | 0.35 | 0.000508 | 0.50 | 0.405358 | 0.48 | 0.239423 | 0.68 | 0.417583 |
| GATA6 | 0.67 | 0.057851 | 0.65 | 1.49E-07 | 0.68 | 0.083787 | 0.98 | 0.965776 | 1.02 | 0.936819 |
| PLK2 | 0.72 | 0.112915 | 0.41 | 2.48E-06 | 0.64 | 0.002456 | 0.58 | 0.142406 | 0.90 | 0.570129 |

**Table S2:** Cardiomyocyte maturity genes

| Gene | Sparse<br>(Dense)<br>FC | p-value | XMU<br>(Dense)<br>FC | p-value | S1P<br>(Dense)<br>FC | p-value | XMU<br>(Sparse)<br>FC | p-value | S1P<br>(Sparse)<br>FC | p-value |
| --- | --- | --- | --- | --- | --- | --- | --- | --- | --- | --- |
| KLF15 | 0.37 | 0.001345 | 0.62 | 0.015381 | 0.40 | 0.081736 | 1.69 | 0.216003 | 1.08 | 0.852327 |
| PPARA | 0.50 | 0.033375 | 0.69 | 0.000397 | 0.51 | 0.059213 | 1.42 | 0.404501 | 1.04 | 0.913521 |
| RB1 | 0.66 | 0.377255 | 1.07 | 0.637022 | 0.80 | 0.801415 | 1.65 | 0.23275 | 1.21 | 0.62234 |
| RXRA | 0.67 | 0.183603 | 0.75 | 0.008464 | 0.66 | 0.090851 | 1.15 | 0.740298 | 1.00 | 0.995685 |
| PPARGC1A | 0.69 | 0.237739 | 0.46 | 2.62E-09 | 0.75 | 0.403987 | 0.69 | 0.305008 | 1.10 | 0.703669 |
| NUPR1 | 0.67 | 0.718043 | 0.03 | 0.00025 | 1.33 | 0.902237 | 0.05 | 0.065356 | 2.04 | 0.421311 |
| PPARD | 0.72 | 0.091847 | 0.88 | 0.254178 | 0.65 | 0.407204 | 1.25 | 0.360958 | 0.90 | 0.70345 |
| STAT5B | 0.96 | 0.925941 | 1.30 | 0.005076 | 0.88 | 0.835451 | 1.38 | 0.238056 | 0.92 | 0.749992 |
| TP53 | 1.04 | 0.919216 | 0.74 | 0.007432 | 1.17 | 0.620798 | 0.73 | 0.178197 | 1.13 | 0.501919 |
| SMARCA4 | 1.38 | 0.318031 | 1.07 | 0.563766 | 1.37 | 0.427076 | 0.79 | 0.484057 | 1.01 | 0.984053 |
| PPARGC1B | 0.57 | 0.118278 | 0.24 | 3.94E-19 | 0.70 | 0.255304 | 0.43 | 0.02966 | 1.24 | 0.451305 |
| SIRT1 | 0.64 | 0.391808 | 1.17 | 0.300566 | 0.70 | 0.460866 | 1.85 | 0.167862 | 1.10 | 0.785253 |
| BRCA1 | 0.70 | 0.444602 | 0.39 | 5.06E-07 | 0.68 | 0.240951 | 0.57 | 0.255112 | 0.97 | 0.929278 |
| ISL1 | 0.70 | 0.72808 | 0.39 | 0.047124 | 1.27 | 0.832355 | 0.57 | 0.575785 | 1.85 | 0.226337 |
| EZH2 | 0.95 | 0.925928 | 0.90 | 0.608155 | 0.99 | 0.989079 | 0.97 | 0.957506 | 1.04 | 0.872184 |
| KLF2 | 1.07 | 0.943608 | 2.03 | 0.307754 | 1.36 | 0.80982 | 1.93 | 0.374266 | 1.31 | 0.606284 |
| TCF7L2 | 1.11 | 0.804331 | 0.82 | 0.208738 | 1.40 | 0.432616 | 0.75 | 0.257579 | 1.28 | 0.274109 |
| IRF3 | 1.22 | 0.675849 | 1.29 | 0.038872 | 1.26 | 0.749532 | 1.08 | 0.891222 | 1.05 | 0.881247 |
| EPAS1 | 0.33 | 1.33E-06 | 0.34 | 1.38E-16 | 0.23 | 0.015726 | 1.04 | 0.945471 | 0.69 | 0.436981 |
| STAT6 | 0.55 | 0.006801 | 0.19 | 2.02E-30 | 0.37 | 0.000199 | 0.35 | 2.87E-05 | 0.68 | 0.125648 |
| HMGA1 | 0.71 | 0.242416 | 0.72 | 0.000562 | 0.74 | 0.087149 | 1.04 | 0.929241 | 1.05 | 0.807651 |
| CREBBP | 0.76 | 0.429739 | 0.98 | 0.906761 | 0.94 | 0.893223 | 1.32 | 0.408709 | 1.24 | 0.3625 |
| RORA | 0.77 | 0.632319 | 1.10 | 0.719185 | 0.81 | 0.681446 | 1.44 | 0.432174 | 1.05 | 0.881326 |
| TBX5 | 0.81 | 0.516316 | 0.73 | 0.015109 | 0.88 | 0.818683 | 0.91 | 0.753349 | 1.09 | 0.682947 |
| HIF1A | 0.91 | 0.827102 | 0.73 | 0.000624 | 0.88 | 0.888031 | 0.82 | 0.57784 | 0.98 | 0.944643 |
| ESRRA | 0.97 | 0.964559 | 0.64 | 0.001142 | 0.66 | 0.201523 | 0.67 | 0.165544 | 0.69 | 0.080643 |
| GATA4 | 0.97 | 0.924665 | 1.54 | 8.22E-09 | 1.08 | 0.894736 | 1.62 | 0.002196 | 1.12 | 0.607182 |
| MEF2C | 1.07 | 0.85549 | 1.26 | 0.043804 | 1.51 | 0.008506 | 1.20 | 0.525246 | 1.42 | 0.054329 |
| MYOCD | 1.22 | 0.587856 | 0.84 | 0.148871 | 1.24 | 0.38708 | 0.70 | 0.236397 | 1.02 | 0.931564 |
| SREBF1 | 0.34 | 0.00046 | 0.32 | 1.42E-11 | 0.40 | 0.00197 | 0.96 | 0.945008 | 1.18 | 0.522472 |
| NR2C2 | 0.54 | 0.352976 | 0.94 | 0.749191 | 0.62 | 0.730363 | 1.77 | 0.374012 | 1.14 | 0.829856 |
| EP300 | 0.70 | 0.467532 | 1.15 | 0.234023 | 0.85 | 0.847678 | 1.66 | 0.220495 | 1.21 | 0.612428 |
| FOXO3 | 0.75 | 0.349721 | 0.86 | 0.130914 | 0.78 | 0.216367 | 1.17 | 0.641334 | 1.05 | 0.805169 |
| CEBPB | 0.94 | 0.922627 | 0.88 | 0.386107 | 1.17 | 0.797417 | 0.96 | 0.952266 | 1.27 | 0.462195 |
| AHR | 0.93 | 0.932517 | 1.35 | 0.172625 | 1.00 | 0.998442 | 1.46 | 0.403747 | 1.07 | 0.848077 |
| ESRRA | 0.97 | 0.964559 | 0.64 | 0.001142 | 0.66 | 0.201523 | 0.67 | 0.165544 | 0.69 | 0.080643 |
| TFAM | 0.98 | 0.972321 | 0.93 | 0.638489 | 1.12 | 0.815031 | 0.96 | 0.925733 | 1.15 | 0.524921 |
| TCF3 | 1.07 | 0.870547 | 1.03 | 0.829302 | 0.99 | 0.976384 | 0.98 | 0.966962 | 0.93 | 0.718576 |
| NR4A3 | 1.11 | 0.880451 | 4.19 | 0.042536 | 2.08 | 0.097711 | 3.84 | 0.020697 | 1.89 | 0.035762 |
| NFATC2 | 0.24 | 8.81E-07 | 0.31 | 4.36E-14 | 0.45 | 0.01823 | 1.33 | 0.590606 | 1.85 | 0.066561 |
| TOB1 | 0.69 | 0.202871 | 0.71 | 0.006508 | 0.64 | 0.047766 | 1.05 | 0.921464 | 0.93 | 0.750685 |
| CTCF | 0.77 | 0.330234 | 0.91 | 0.388474 | 0.73 | 0.337073 | 1.20 | 0.520739 | 0.95 | 0.79525 |
| HDAC1 | 0.93 | 0.868417 | 1.14 | 0.307232 | 0.80 | 0.595919 | 1.25 | 0.423766 | 0.86 | 0.476421 |
| E2F6 | 0.98 | 0.964822 | 1.04 | 0.850897 | 0.98 | 0.95372 | 1.08 | 0.834633 | 1.00 | 0.980775 |
| GMNN | 1.01 | 0.983859 | 0.77 | 0.075827 | 1.04 | 0.948911 | 0.78 | 0.420264 | 1.04 | 0.882408 |
| USF1 | 1.19 | 0.652289 | 1.01 | 0.9356 | 1.18 | 0.807159 | 0.87 | 0.699954 | 1.01 | 0.985944 |
| STAT5A | 1.44 | 0.745261 | 0.72 | 0.622383 | 0.71 | 0.869697 | 0.51 | 0.548587 | 0.49 | 0.406925 |
| ATF3 | 1.39 | 0.496897 | 6.55 | 0.001271 | 4.69 | 3.45E-13 | 4.80 | 0.001356 | 3.44 | 3.16E-05 |

**Table S3:**
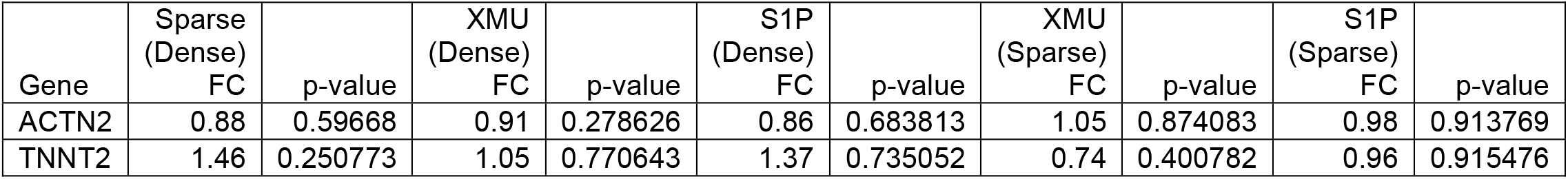
Cardiomyocyte-specific genes

